# Wide-field Intensity Fluctuation Imaging

**DOI:** 10.1101/2023.07.29.551117

**Authors:** Qingwei Fang, Alankrit Tomar, Andrew K. Dunn

## Abstract

The temporal intensity fluctuations contain important information about the light source and light-medium interaction and are typically characterized by the intensity autocorrelation function, *g*_2_(*τ*). The measurement of *g*_2_(*τ*) is a central topic in many optical sensing applications, ranging from stellar intensity interferometer in astrophysics, to fluorescence correlation spectroscopy in biomedical sciences and blood flow measurement with dynamic light scattering. Currently, *g*_2_(*τ*) at a single point is readily accessible through high-frequency sampling of the intensity signal. However, two-dimensional widefield measurement of *g*_2_(*τ*) is still limited by camera frame rates. We propose and demonstrate a 2-pulse within-exposure modulation approach to break through the camera frame rate limit and obtain the quasi *g*_2_(*τ*) map in wide field with cameras of only ordinary frame rates.

**Highlights:**

- The quasi intensity autocorrelation function can be measured at almost arbitrary time lags
- Cameras of ordinary frame rates are sufficient for the measurement
- Illumination within the camera exposure is modulated by an acoustic optical modulator using two short pulses

## 1. Introduction

The intensity correlation function is widely used for quantifying optical fluctuations and its measurement has great physical and physiological significance in many optical sensing applications. It was first introduced in 1956 as intensity interferometry to measure the apparent angular diameter of stars^1^ and recently used to study stellar emission processes and calibrate star distances in astrophysics^2,3^. It plays an essential role in fluorescence correlation spectroscopy (FCS) in determining the diffusion coefficient of molecules and investigating biomolecular interaction processes^4–8^. It is used to achieve super-resolution optical fluctuation imaging (SOFI)^9–11^. It is used for particle sizing in dynamic light scattering^12–14^. In addition, tissue blood flow and perfusion information can be extracted from it, on which the diffuse correlation spectroscopy^15–17^(DCS) and laser speckle contrast imaging^18–22^ (LSCI) are developed.

Generally, the intensity autocorrelation function measures the similarity of the intensity signal with itself between now and a moment later. Technically, it is defined as 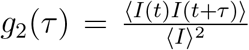 where *τ* is called the time lag, *I*(*t*) is the intensity signal of interest and ⟨ ⟩ denotes averaging. To resolve the intensity autocorrelation function, high temporal resolution detectors, such as avalanche photo-diodes (APD) or single photon avalanche diodes (SPAD), are required to record the intensity at sufficiently high sampling rates. However, these one-dimensional detectors are only capable of single-point *g*_2_(*τ*) measurements.

Light sheet or total internal reflection microscopy enables 2-dimensional (2D) FCS measurements at thousands of locations simultaneously by camera-based fluorescence intensity recording^23,24^. However, the electron multiplying charge coupled devices (EM-CCD), which are currently considered as the most suitable option in comprehensive comparison with other types of 2D detectors^25,26^, are still limited in frame rates (∼1000 frames per second) and suffer from high instrumentation cost.

In DCS and LSCI, some high-speed cameras have been used to record the raw laser speckle signal and measure *g*_2_(*τ*) in a 2D field of view (FOV)^27,28^. In addition, SPAD arrays are utilized by speckle contrast optical spectroscopy to create synthetic multiple-exposure speckle contrast data^29–31^. However, both methods suffer from limited field of view and high instrumentation cost to resolve the signal at sufficient frame rates. Recently rolling shutters have been demonstrated helpful in alleviating the frame rate limit of cameras^32^. But the method trades the spatial resolution for temporal resolution and cannot measure *g*_2_(*τ*) of slowly varying dynamics due to the limited length of elongated speckle patterns created by the elliptical aperture.

Overall, *g*_2_(*τ*) is still measured based on the fully time-resolved signal, from which, however, high-frequency signal sampling is inevitable. As such, current methods must sacrifice either field of view or spatial resolution to accelerate the signal sampling. Here we propose a method to measure *g*_2_(*τ*) without resolving the fast temporal dynamics of the signal, thereby enabling characterization of rapid intensity fluctuations even at low camera frame rates.

Our method borrows the idea of speckle contrast from LSCI. The relationship between speckle contrast *K* and *g*_2_(*τ*) has been well established in LSCI, where the pixel intensity *S*(*T*) is defined as

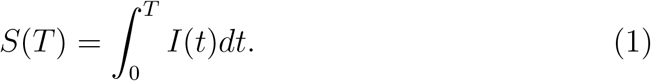

The speckle contrast is then defined as

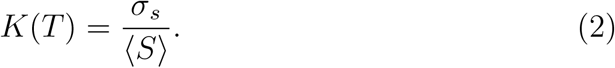

Speckle contrast can be calculated either spatially or temporally. Spatially, a *N* × *N* sliding window is typically used across the image to generate the speckle contrast of the center pixel by computing the standard deviation and mean of all *N*^2^ pixel intensities within the window under the assumption of ergodicity^21^. Temporally, a series of images with the same camera exposure time can be acquired to calculate the speckle contrast at a certain pixel by computing the standard deviation over the mean of the pixel’s intensity in those images.

Note that speckle contrast can be measured with a much lower frame rate than *g*_2_(*τ*) since it is based on statistical properties of the *integrated* signal, *S* within the camera exposure time *T*. The integrated signal’s speckle contrast *K* within different *T* can be measured by multiple exposures of different exposure times^33–36^. The camera exposures do not have to be consecutive or acquired with a fast frame rate as long as the statistical property of the signal remains unchanged over the multiple exposures.

Speckle contrast is related to *g*_2_(*τ*) in a way that *K*^2^ is an integral of *g*_2_(*τ*) weighted by a right triangle function (*T* − *τ*) if the illumination is held constant within the camera exposure^37^

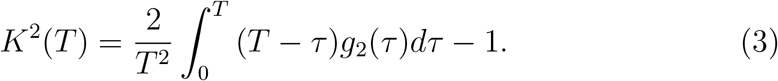

Recently, Siket et al. have generalized the relationship to cases where the illumination can be modulated by an arbitrary waveform^38^ (Supplemental section S1), namely

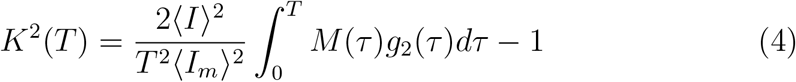

where *I*_*m*_ is the modulated signal intensity and *I*_*m*_(*t*) = *I*(*t*)*m*(*t*) where *m*(*t*) is the modulation waveform within the camera exposure and ranges from 0 to 1. *M* (*τ*) is the autocorrelation of the modulation waveform defined as 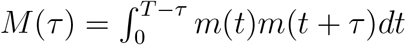.

One important observation from Eq. 4 is that if we could find a modulation waveform *m*(*t*) such that *K*^2^(*T*) = *g*_2_(*T*), then we can measure *g*_2_(*τ* = *T*) by measuring *K*^2^(*T*) at a much lower frame rate. We achieve this by 2-pulse modulated multiple-exposure imaging with two illumination pulses placed inside the exposure and their temporal separation (denoted as *T*) varied across exposures (Fig. 1a). The laser illumination pulses are created by externally modulating the laser with an acoustic optical modulator (AOM). The idea is that when the pulse duration *T*_*m*_ is approaching 0, *M* (*τ*) weighted by 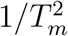 would become the sum of two delta functions (Fig. 1b) and Eq. 4 would be reduced to

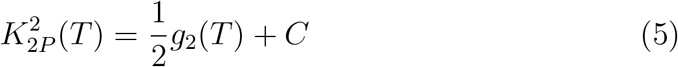

where 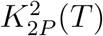 represents the square of the 2-pulse modulated speckle contrast and *C* is a constant that is independent of *T* (Methods 4.1). Since 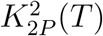 forms a linear relationship with *g*_2_(*T*), we call it quasi *g*_2_(*τ*).

**Figure 1.**
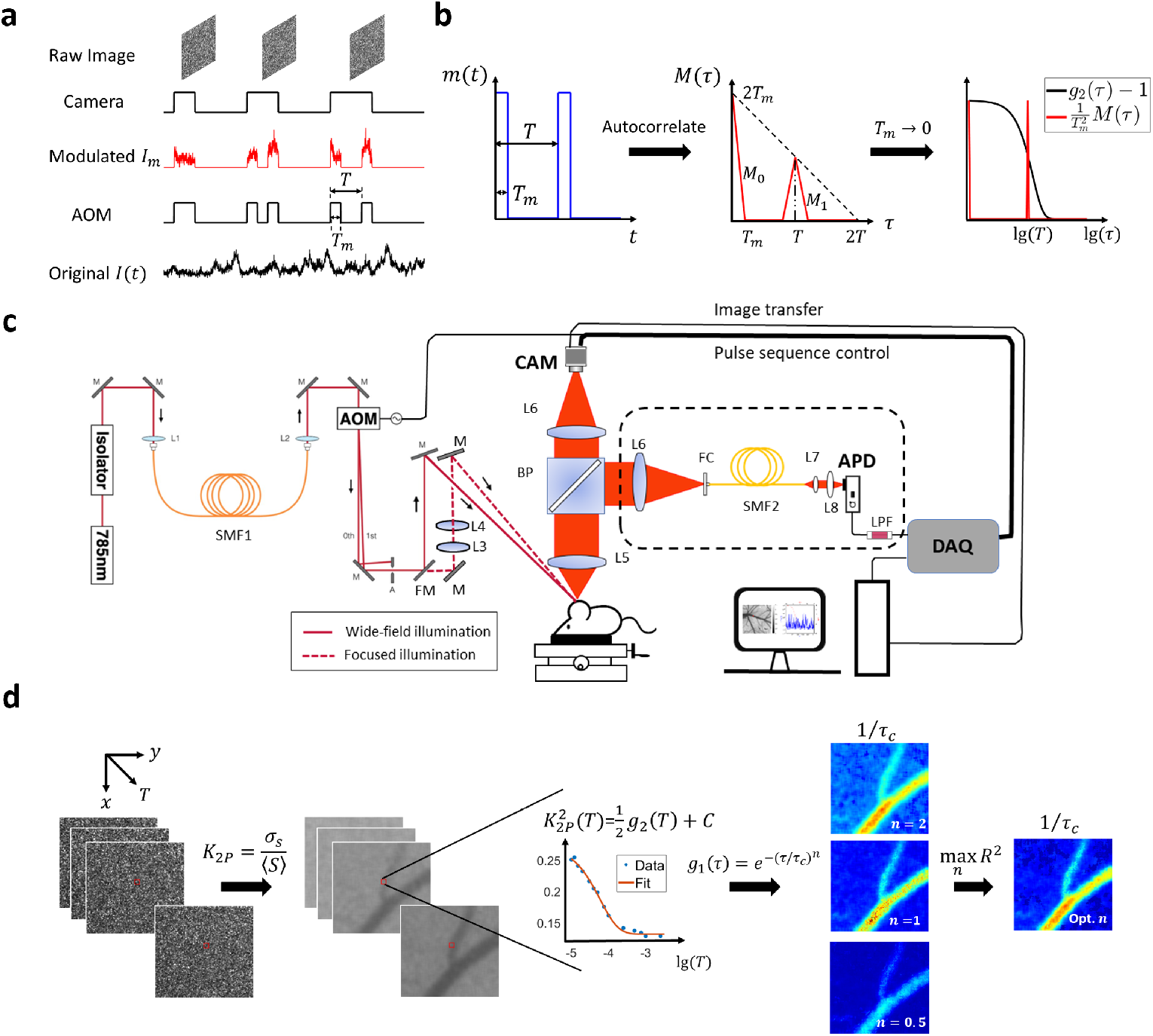
Overview of the methodology and instrumentation. **a** Temporal relationship between intensity modulation and camera exposure. The *x*-axis is time. AOM: acoustic optical modulator. The sample is illuminated only when AOM modulation voltage is high. Hence for *I*_*t*_, only the signal when AOM is high will be recorded and integrated onto the camera raw image. *T* : the period of the two-pulse modulation waveform. *T*_*m*_: pulse duration. **b** The autocorrelation function of 2-pulse modulation waveform. *m*(*t*): intensity modulation waveform. *m*(*t*) ∈ [0, 1]. *M* (*τ*): the autocorrelation of *m*(*t*). 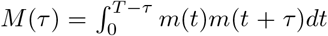. *M* (*τ*) consists of two pulses denoted as *M* and *M*. When *T*_*m*_ is approaching 0, 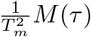 becomes the sum of two delta functions. **c** Diagram of the instrumentation of this study. Two illumination light paths are constructed: widefield (real line) and focused (dashed line). The light is switched between the two paths by a flip mirror but modulated by the same pulse sequence. The back-scattered light is collected by the objective and then split by a 50/50 beamsplitter. The two splits are collected by camera and APD, respectively. AOM: acoustic optical modulator. BP: 50/50 beamsplitter plate. APD: avalanche photon diode. DAQ: data acquisition board. CAM: camera. L: lens. M: mirror. FM: flip mirror. FC: fiber coupler. SMF: single-mode fiber. LPF: low-pass filter. **d** The workflow of extracting correlation time from 2-pulse modulated multiple-exposure raw images. The 2-pulse modulated specke contrast, *K*_2*P*_, is first computed from the modulated raw speckle images and its trace along the third dimension *T* is then fitted with different electric field correlation *g*_1_(*τ*) models (*n* = 2, 1 or 0.5). The best *g*_1_(*τ*) model is identified by maximizing the coefficient of determination, *R*^2^. lg: logarithm to base 10.

Cameras of ordinary frame rates as low as 1 Hz are sufficient for our *g*_2_(*τ*) measurement approach as long as the signal’s statistical property remains invariant within the measurement. The camera-based characterization of *g*_2_(*τ*) at various time lags is first validated against the *g*_2_(*τ*) curve obtained by the traditional single-point photodiode measurement with 1 MHz sampling rate (instrumentation shown in Fig. 1c). Furthermore, wide-field quasi *g*_2_(*τ*) measurement and correlation time mapping are demonstrated in case of *in vivo* blood flow imaging (workflow summarized in Fig. 1d).

## 2. Results

### 2.1 A 10 μs pulse duration is short enough such that 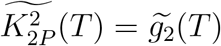

In this section, we first verify the equivalency between normalized 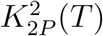 and *g*_2_(*T*) (denoted as 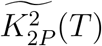 and 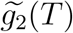, respectively) with numerical simulation using a *T*_*m*_ = 10 *μ*s pulse duration. As highlighted by the green dashed line in Fig. 2a, the 2-pulse modulated 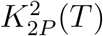 decreases to the flat level earlier than the unmodulated *K*^2^(*T*) but at about the same time as *g*_2_(*τ*). The shape of 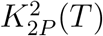 curve is also more similar to that of *g*_2_(*τ*) compared with *K*^2^(*T*). Further normalization reveals the consistency between the normalized 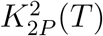 and *g*_2_(*τ*) curves (Fig. 2b). Such equivalency holds for *g*_2_(*τ*) curves over a wide range of decreasing speeds. The correlation time *τ*_*c*_ is varied from 5 *μ*s to 0.1s, which covers the whole spectrum of *τ*_*c*_ reported by Postnov et al^27^. The discrepancy between 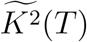 and 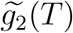 drastically increases when *τ*_*c*_ is reduced to 5 *μ*s (Fig. 2c, d). However, 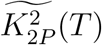 maintains a good consistency with 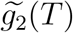 throughout the *τ*_*c*_ range (Fig. 2c, d). The maximum relative percentage discrepancy is below 10^−5^%.

**Figure 2.**
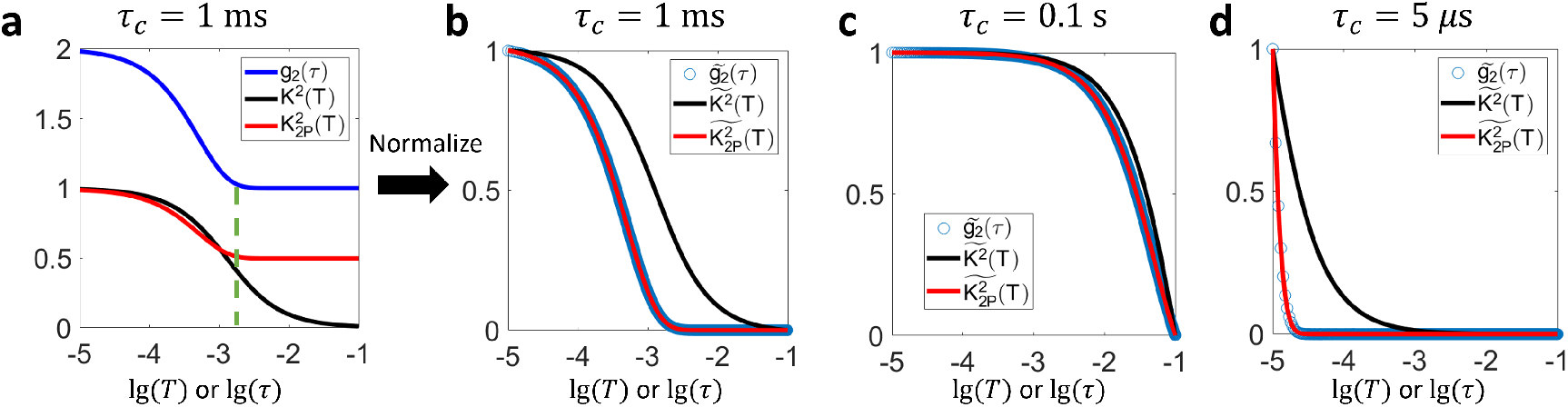
Numerical simulation of *K*^2^ and 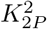 curves given the same *g*_2_(*τ*). **a** Comparison of the given *g*_2_(*τ*) curve with speckle contrast curves with and without 2-pulse modulation. 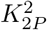 denotes the square of 2-pulse modulated speckle contrast while *K*^2^ represents the square of speckle contrast without within-exposure modulation. The pulse duration in 2-pulse modulation is *T*_*m*_ = 10 *μ*s. *τ*_*c*_ = 1 ms. **b** Comparison of the three curves after separate normalization on the *y*-axis so that the dynamic range is all normalized to [0, 1]. **c** Comparison of normalized *g*_2_(*τ*), *K*^2^(*T*) and 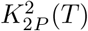 when *τ*_*c*_ = 0.1 s. **d** Comparison of normalized *g*_2_(*τ*), *K*^2^(*T*) and 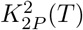 when *τ*_*c*_ = 5 *μ*s. lg represents the logarithm to base 10 throughout this paper.

The consistency between normalized 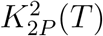 and *g*_2_(*T*) with a 10 *μ*s pulse duration holds experimentally as well. For *in vitro* microfluidics experiments, raw images of different exposure times under 2-pulse modulation are shown in Fig. 3a. The average pixel intensity is approximately the same across different camera exposures, which is expected since the effective exposure time is kept the same in those exposures, i.e. all 20 *μ*s. But the corresponding speckle contrast shows significant decrease when the temporal separation between the two illumination pulses, *T*, increases from 50 to 100 *μ*s (Fig. 2b). Such trend is further illustrated in Fig. 3c where the 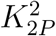 in the microfluidic channel area decreases as *T* increases. In addition, 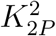 of higher flow rates begins to decrease earlier than those of lower rates. Such relationship between flow rate and start time of decreasing is also reflected in *g*_2_(*τ*) curves which are derived from APD measurements of 1 MHz sampling rate (Fig. 3d). Most importantly, When 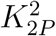 and *g*_2_(*τ*) curves are normalized, they overlap on each other (Fig. 3e).

**Figure 3.**
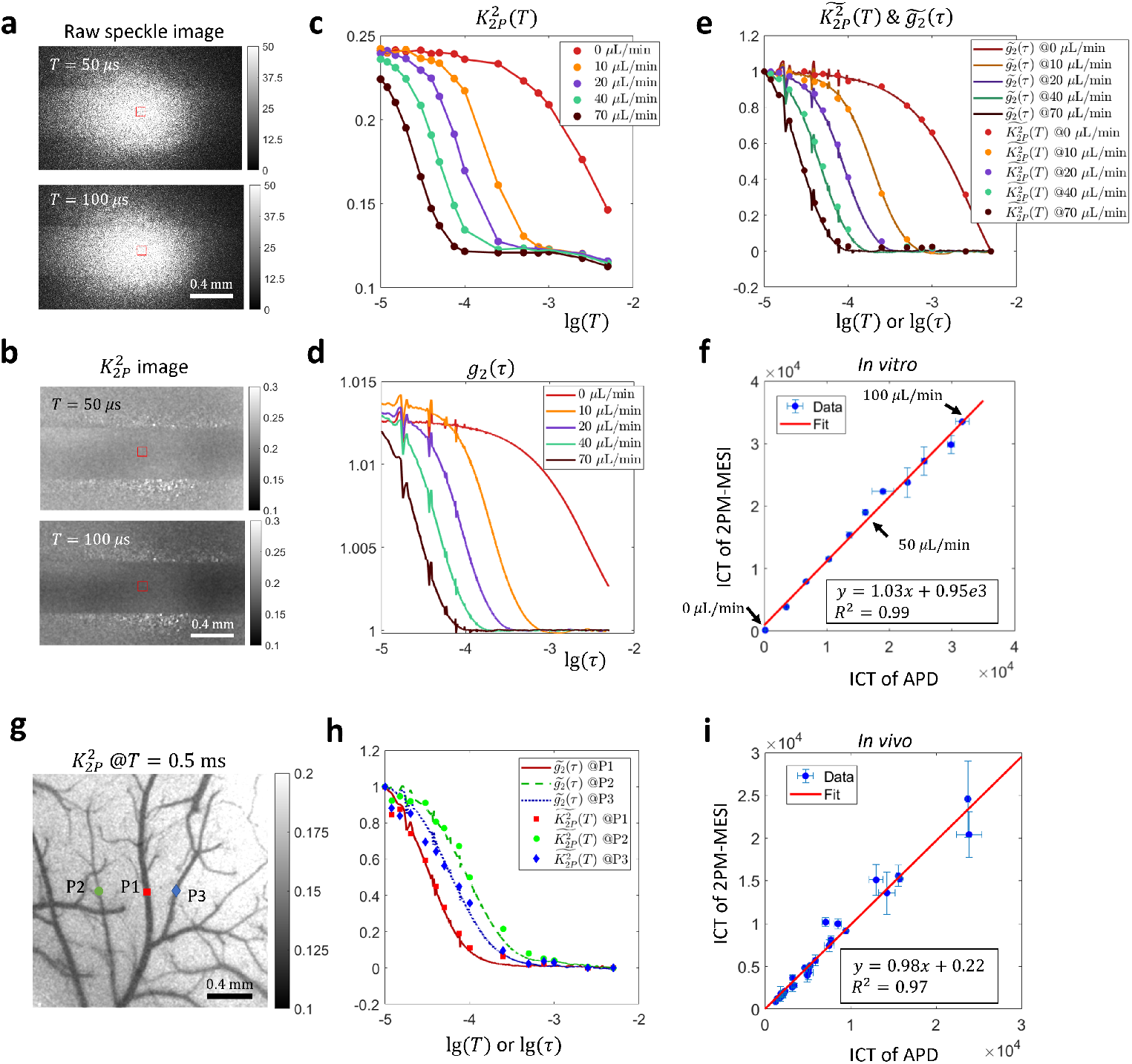
Experimental validation of the consistency between normalized 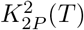 and *g*_2_(*τ*) curves *in vitro* and *in vivo* under focused illumination. **a**-**f** *In vitro* microfluidic experiment results. **g**-**i** *In vivo* experiment results. **a** Raw images acquired in 2-pulse modulation strategy. **b** Speckle contrast images calculated from 2-pulse modulated raw images. **c** The 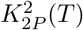 curves extracted from the microfluidic channel region (ROI shown by red boxes in a, b) in various flow rates. **d** The *g*_2_(*τ*) curves in the channel region in various flow rates. **e** Comparison of normalized *g*_2_(*τ*) and normalized 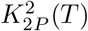. **f** Comparison of ICT values extracted from *g*_2_(*τ*) and 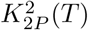 curves. Data points from 11 flow rates ranging from 0 to 100 *μ*L/min with a step size of 10 *μ*L/min are shown. The optimal *g*_1_(*τ*) model is first identified by the fitting algorithm through maximizing *R*^2^, the coefficient of determination, for both camera and APD measurements assuming three different *g*_1_(*τ*) models, i.e. 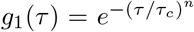 and *n* = 2, 1, or 0.5. The ICT of the optimal *g*_1_(*τ*) model is then used for comparison. **g** Position of three representative points in the FOV *in vivo* where APD measurements are performed. The background 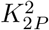 image is acquired under widefield illumination. **h** Comparison of normalized *g*_2_(*τ*) and 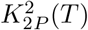 curves at the three points. **i** Comparison of ICT values extracted from *g*_2_(*τ*) and 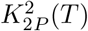 curves *in vivo*. The *n* = 1 *g*_1_(*τ*) model is used for curve fitting. 28 points from 4 mice are shown.

The downtick in the tail of 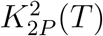 curves (Fig. 3c), which is not present in corresponding *g*_2_(*τ*) curves (Fig. 3d), arises from the incomplete gating of light by AOM between the two illumination pulses. When the distance between two illumination pulses, *T*, becomes too large relative to the pulse duration, the effects of non-zero residual illumination accumulated in between are no longer negligible and can result in a lowered 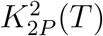 value (Supplemental Material section S3). Removing the downtick will be the topic of a future publication.

Apart from comparing the values of normalized 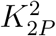 and *g*_2_(*τ*), we also compared the inverse correlation time (ICT), i.e. 1*/τ*_*c*_, extracted from 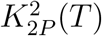 and *g*_2_(*τ*) curves. First, the electric field autocorrelation *g*_1_(*τ*) model identification capability of 2-pulse modulated multiple-exposure speckle imaging (2PM-MESI) is validated against APD-based direct *g*_2_(*τ*) measurements. When the flow is zero, for the best *g*_1_(*τ*) model, *n* = 1. When the flow is non-zero among the tested flow rates, *n* = 2 (Supplemental Fig. S2). Second, as expected, the larger the flow rate, the larger the ICT (Fig. 3f). Third, ICT extracted from 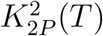 curves consisting of only 15 values of *T* is consistent with that from *g*_2_(*τ*) evaluated at a much denser set of *τ* (*R*^2^ = 0.99, Fig. 3f). This suggests that there is redundancy in *g*_2_(*τ*) curves and that the correlation time of *g*_2_(*τ*) can be estimated from only a few key data points.

The 2-pulse modulated 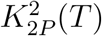 curve is also compared with *g*_2_(*τ*) *in vivo*. Fig. 3g shows the location of three points (P1-3) where single-point direct *g*_2_(*τ*) measurements are performed with an APD. Note that their vessel radii are different, i.e., P1 the largest, P2 the smallest and P3 in the middle. As expected, their *g*_2_(*τ*) curves are also separated, i.e. *g*_2_(*τ*) of P1 starts decreasing first while that of P2 does last (Fig. 3h). The normalized *in vivo* 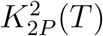 curve is not as consistent with that of *g*_2_(*τ*) as it is *in vitro*. It could be due to the stronger flow disturbance *in vivo* as evident with the large error bars in the ICT plot (Fig. 3i). Note that APD and 2PM-MESI measurements are not performed simultaneously since 2PM-MESI requires modulating the illumination while the other not. The better consistency between normalized 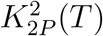 and *g*_2_(*τ*) *in vitro* could arise from the better flow control *in vitro*.

The *g*_1_(*τ*) model identification capability is also degraded *in vivo*. ICT extracted from 2PM-MESI 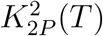 curves is consistent with that from *g*_2_(*τ*) curves measured with APD when the *g*_1_(*τ*) model is fixed to *n* = 1 for both APD and 2PM-MESI measurements (Fig. 3i, *R*^2^ = 0.97). Unfixing the model and let the algorithms choose the optimal *n* based on the fitting performance results in a degraded consistency of ICT between APD and 2PM-MESI measurements (Supplemental Fig. S3, *R*^2^ = 0.94). It indicates that for complex flow dynamics *in vivo*, there is still room for the current settings of *T* of 2PM-MESI, e.g. the number of exposures and values of *T*, to be further optimized. In addition, a single *g*_1_(*τ*) model might be insufficient *in vivo* and a mixed model might be warranted^27^.

### 2.2 Widefield Quasi g_2_(τ) Measurement and Correlation Time Mapping

The correlation time is an important indicator of blood flow speed in LSCI. To demonstrate the 2D quasi *g*_2_(*τ*) measurement and correlation time mapping capability of our method, we present in this section widefield illumination results. APD results are not shown because they are dominated by noise in this illumination regime. Fig. 4a-d show the 2-pulse modulated quasi *g*_2_(*τ*) images in wide-field illumination at various *τ* = *T*. Small vessels gradually appear as *T* increases, indicating slower intensity fluctuations. Note that the image size is as large as 1000×750 pixels (the corresponding FOV under 2× magnification: ∼ 2.9 × 2.2 mm^2^). Fig. 4e-g show the inverse correlation time (ICT) maps extracted with the three different *g*_1_(*τ*) models. It can be seen that the ICT map of optimal *n* (Fig. 4h) preserves the high ICT values in vascular regions in *n* = 1 and *n* = 2 ICT maps (Fig. 4e, f) as well as the low ICT values in parenchyma regions in *n* = 0.5 ICT map (Fig. 4g). In addition, the distribution of optimal *n* across the field of view (Fig. 4i) is consistent with what is reported by Postnov and Liu et al measuring *g*_2_(*τ*) with high-speed cameras^27,39^. The fitting results of 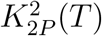 curves at three representative points are shown in Fig. 4j-l (*n* = 2, 1 and 0.5, respectively, position shown in Fig. 4i). The fitting results indicate that 2PM-MESI is capable of identifying one proper *g*_1_(*τ*) model according to the measured quasi *g*_2_(*τ*) curve.

**Figure 4.**
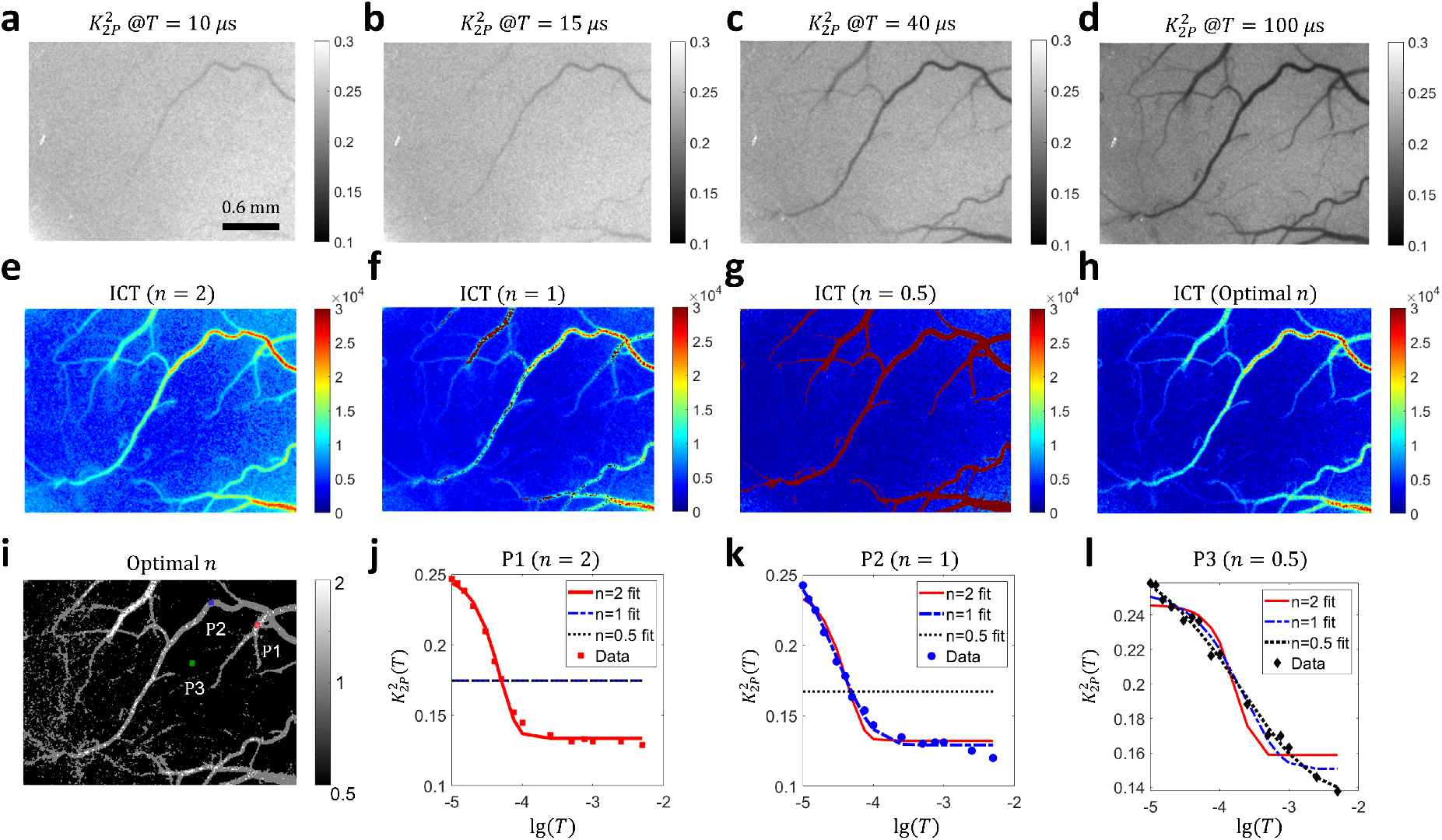
2-pulse illumination modulation across the entire field of view enables wide-field quasi *g*_2_(*τ*) measurement and correlation time mapping. **a**-**d** The quasi *g*_2_(*τ*), i.e. 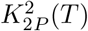 images at *T* = 10 *μ*s, 15 *μ*s, 40 *μ*s and 100 *μ*s, respectively. Image size: 1000 × 750. **e**-**f** ICT maps extracted with three 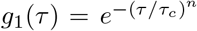 models. *n*=2, 1 and 0.5, respectively. ICT=1*/τ*_*c*_. The 2D map of correlation times was obtained by fitting 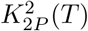 maps at 15 *T* time points ranging from 10 *μ*s to 5 ms. **h** ICT map with *n* optimized at each pixel to maximize the *R*^2^, the coefficient of determination. **i** Map of optimal *n*. **j**-**l** Fitting results of 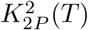 at the three points highlighted in **i**. lg: logarithm to base 10. In figure **j**, the *n* = 1 and *n* = 0.5 *g*_1_(*τ*) models fail to fit the 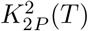 curve. Hence, they appear as a flat line in the plot. The same is true for the *n* = 0.5 *g*_1_(*τ*) model in figure **k**.

## 3. Discussion

The measurement of intensity autocorrelation function, *g*_2_(*τ*) is a fundamental tool in many optical sensing applications to quantify intensity fluctuations and thus investigate the light source and light-medium interaction. However, *g*_2_(*τ*) measurements at short time lags are limited by the camera frame rate. To properly sample the rapid dynamics of intensity fluctuations, traditional two-dimensional *g*_2_(*τ*) measurement methods must sacrifice either field of view or spatial resolution to increase the temporal sampling rate. We propose the 2-pulse within-exposure modulation approach to break through the camera frame limit and change the problem from fast acquisition of raw images to the fast modulation of laser illumination. We showed that the normalized *g*_2_(*τ*) can be well approximated by the normalized 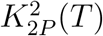, the 2-pulse modulated speckle contrast. With our method, *g*_2_(*τ*) can be measured at short time lags independent of camera frame rate.

The smallest time lag at which *g*_2_(*τ*) can be characterized by the 2-pulse modulated multiple-exposure imaging depends on the smallest value of *T* that can be achieved. Since *T* must be greater than or equal to the pulse duration *T*_*m*_, the question becomes how short the illumination pulse could be made while achieving a sufficient signal-to-noise ratio. We demonstrated that with a 10 *μ*s pulse duration with ∼100 mW laser power input into the AOM, the quasi *g*_2_(*τ*) can be measured with a decent signal-to-noise ratio even in widefield illumination, which is already beyond the capability of most cameras to measure *g*_2_(*τ*) with the traditional method. With a 10 *μ*s pulse duration, we are able to evaluate quasi *g*_2_(*τ*) at the smallest time lag of *τ* = 10 *μ*s, which would otherwise require a camera frame rate of 100 kHz with traditional methods.

Theoretically, the measurement of *g*_2_(*τ*) can be made at almost arbitrary time lags except those smaller than the pulse width *T*_*m*_. The sampling of the *g*_2_(*τ*) curve is determined by the number and values of the time lags. Finer sampling of the *g*_2_(*τ*) curve requires more images to be acquired, but is still independent of camera frame rate. The advantage of 2-pulse modulated multiple-exposure imaging over the traditional *g*_2_(*τ*) measurement method is that it enables cameras of even ordinary frame rates to measure *g*_2_(*τ*) at user-specified time lags. Even though it requires the use of an AOM or similar gating hardware to modulate the illumination within the camera exposure, the overall instrumentation cost is still substantially lower than that of high-speed cameras. In addition, the use of pulsed illumination reduces the average power incident upon the sample compared with continuous illumination, which can reduce tissue damage or photo-bleaching of fluorophores.

Note that we perform the modulation of the illumination, but our method is not limited to this case, especially in applications where modulation in the signal detection end is more convenient, for example intensity interferometry. In this case, modulation happens after the light interacts with the medium. Modulation could be also applied in the signal post-processing phase instead of the imaging phase, which we will revisit soon in the following discussion.

Even though the 2-pulse modulation strategy borrows the idea of speckle contrast from LSCI, it has the potential of being generalized to other optical applications than LSCI. This is because speckle contrast is, in definition, identical to the variation coefficient of a general signal. The relationship between speckle contrast and *g*_2_(*τ*) holds without special properties that would distinguish speckle from other types of intensity signal (Supplemental sections S1 and S2). Therefore, the idea of approximating *g*_2_(*τ*) with speckle contrast is not limited to speckle intensity signals. We verify the hypothesis with fluorescent intensity signal. We confirm that the 2-pulse modulation strategy works in FCS as well in principle by applying 2-pulse modulation to a realistic fluorescence intensity signal in the signal post-processing phase (Fig. 5).

**Figure 5.**
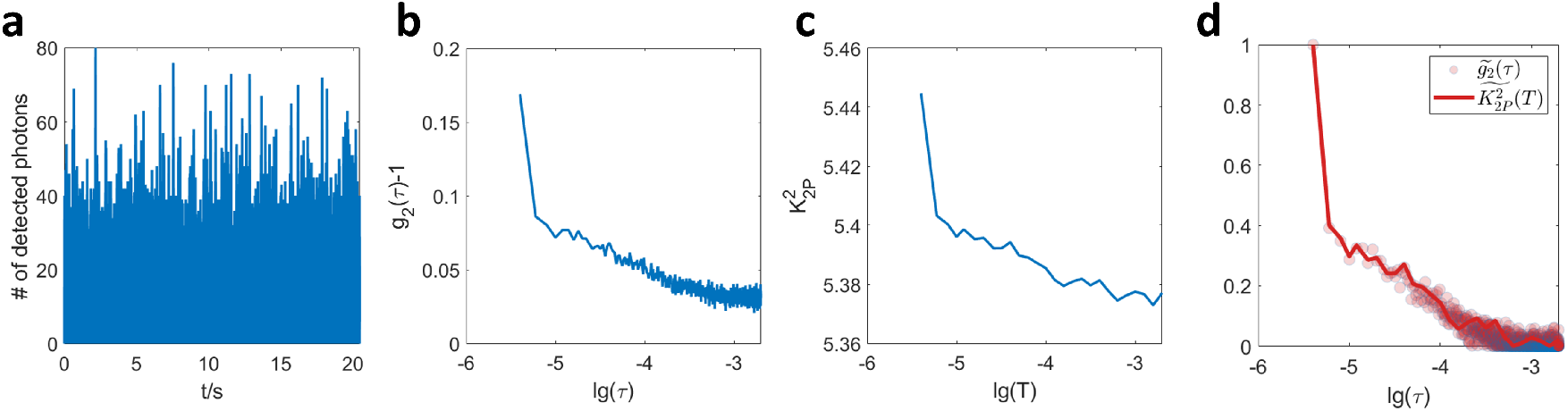
Proof of concept that the 2-pulse modulation strategy works in fluorescence correlation spectroscopy as well. **a** The photon count vs. time plot of a 33 nM Cy5 dye solution recorded by a confocal microscope. Sampling rate: 500 kHz. **b** The plot of *g*_2_(*τ*) curve. **c** The plot of 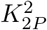 curve. When calculating 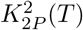, the pixel intensity is generated by gating and integrating the fluorescent intensity signal. Pulse width *T*_*m*_ = 2 *μ*s. **d** Comparison of normalized *g*_2_(*τ*) and 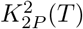 curves for the fluorescent intensity signal.

We have demonstrated the relative equivalency between *g*_2_(*τ*) and 2-pulse modulated speckle contrast, i.e. 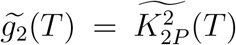. This is enough if we only care about the correlation time of *g*_2_(*τ*) since correlation time is invariant to linear transformations of *g*_2_(*τ*). However, sometimes the absolute value of *g*_2_(*τ*) matters. Does our method still work in this case? We look into this question through applying 2-pulse modulation in the signal post-processing phase again. According to Eq. 5, the absolute value of *g*_2_(*τ*) can be estimated from 2-pulse modulated speckle contrast 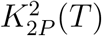, i.e. 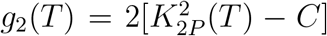. As shown in Fig. 6, the *g*_2_(*τ*) curves of realistic APD signals and the 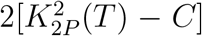 curves calculated from the same APD signal match with each other absolutely even though not perfectly. It suggests that given the same signal collected by exactly the same instrumentation, the absolute value of *g*_2_(*τ*) of the signal can be estimated from its 2-pulse modulated speckle contrast. However, to recover the absolute values of *g*_2_(*τ*) accurately, the requirement on the pulse duration is higher than to just recover the relative values (Supplemental section S4).

**Figure 6.**
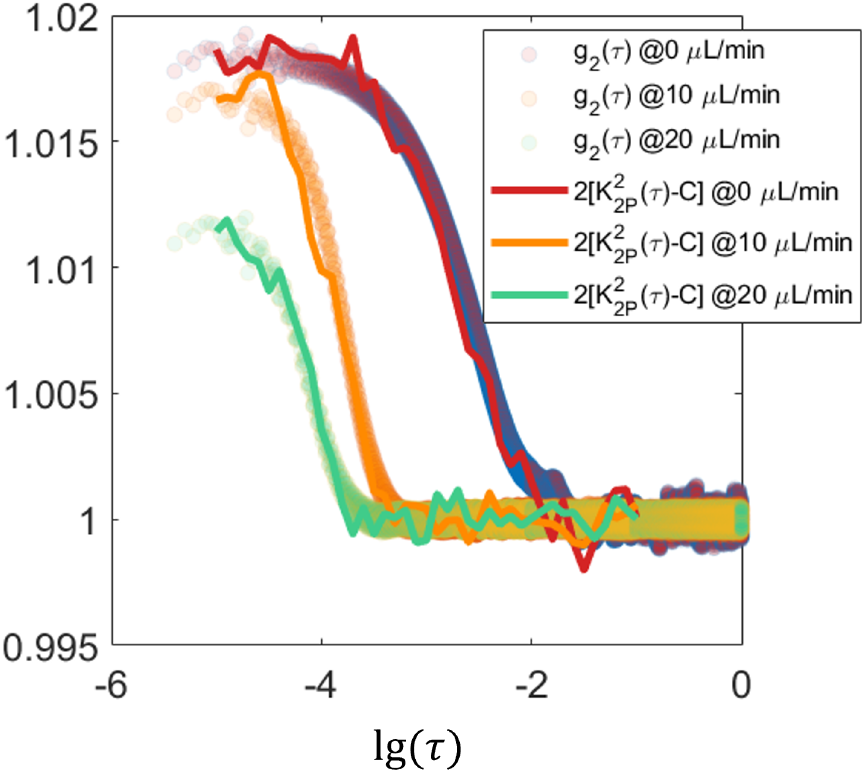
Evaluating the absolute estimation of *g*_2_(*τ*) based on 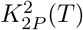 through applying 2-pulse modulation in the signal post-processing phase over speckle intensity recordings. The speckle signal was acquired by APD on microfluidic devices. Temporal 2-pulse modulated speckle contrast is calculated from pixel intensities generated by gating and summing the realistic APD recordings of speckle signal.

In summary, we proposed the 2-pulse modulation waveform to address the question of measuring *g*_2_(*τ*) without resolving the fast temporal dynamics of the intensity signal of interest and demonstrated wide-field intensity fluctuation imaging. Under the 2-pulse modulation, the problem is essentially converted from how fast the raw intensity images can be acquired to how fast the laser illumination or the detected signal can be modulated within the camera exposure. With a short pulse duration, the normalized *g*_2_(*T*) can be approximated by the normalized 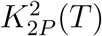. The multiple exposures to acquire 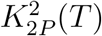 at different *T* do not need to be consecutive or acquired with a fast frame rate. It allows cameras of even ordinary frame rates to characterize the decay of intensity autocorrelation function. The method is expected to enable the 2-dimensional measurement of quasi *g*_2_(*τ*) and facilitate extracting the correlation time in wide field with a substantially lower instrumentation cost.

## 4. Method

### 4.1 Theory

In this section, we explain why 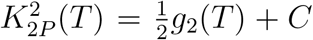 is true when the pulse duration is approaching 0 in 2-pulse modulation. For a 2-pulse modulation waveform (Fig. 1b), *m*(*t*) can be written as

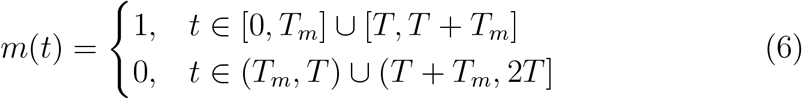

where *T*_*m*_ is the duration of a single illumination pulse and *T* is the period of the modulation waveform. The duty cycle is *d* = *T*_*m*_*/T*. In this case, Eq. 4 can be simplified to

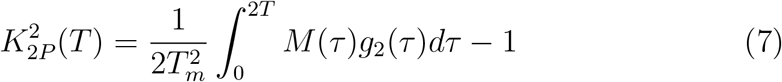

where the subscript 2*P* denotes the speckle contrast under 2-pulse modulation. The corresponding autocorrelation function of the modulation waveform, *M* (*τ*), is a pulse train consisting of two triangle pulses *M*_0_(*τ*) and *M*_1_(*τ*) (Fig. 1b),

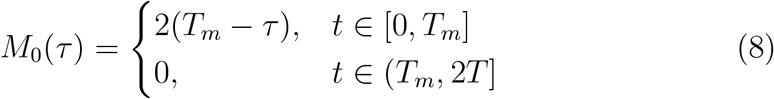

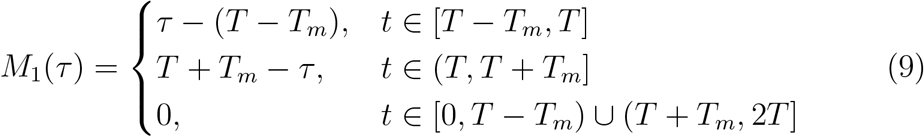

*M* (*τ*) = *M*_0_(*τ*) + *M*_1_(*τ*) and is valid for either *d <*= 0.5 or *d >* 0.5.

Note that the first right triangle pulse *M*_0_(*τ*) is solely dependent on *T*_*m*_ and independent of the temporal separation of the two illumination pulses *T*. In addition, the shape of *M*_1_(*τ*), specifically the width and height, is independent of *T* too. The horizontal position of *M*_1_(*τ*), however depends on *T*. Therefore, when *T* is varied while holding *T*_*m*_ constant, the second triangle pulse *M*_1_(*τ*) will move horizontally. Since 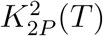 is the integral of the product of *M* (*τ*) and *g*_2_(*τ*), the second triangle pulse, *M*_1_(*τ*), sweeps the *g*_2_(*τ*) curve as *T* changes, through which selective sampling of the *g*_2_(*τ*) curve is achieved.

Scaling *M* (*τ*) by 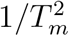 as in Eq. 7, we find that the height/width ratio of its two triangle pulses increases as the illumination pulse duration *T*_*m*_ decreases. In the special case where *T*_*m*_ approaches 0, *M* (*τ*) weighted by 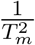 becomes the sum of two delta functions

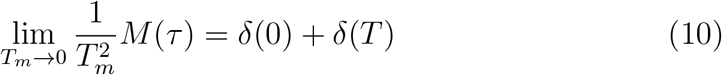

Therefore, Eq. 7 simplifies to Eq. 5 where *C* is a constant representing the contribution of *M*_0_(*τ*) which is independent of *T*. Specifically, 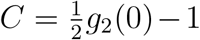 when the impact of *T*_*m*_ on *C* is negligible. A rigorous proof of Eq. 5 in this case from the perspective of statistics is provided in Supplemental section S2. Note that this supplemental proof is valid without assuming Eq. 3, 4 or 7. When *T*_*m*_ must be accounted for, 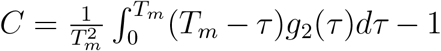 can also be estimated from the 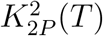 curve since 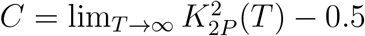 if *g*_2_(*τ*) decreases to 1 when *τ* is infinitely large. Removing the constant *C* and the scaling factor 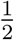 in Eq. 5 through normalization, we have

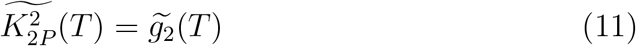

where 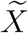 represents the normalization of *X*, i.e. 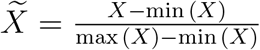.

### 4.2 Numerical simulation

For any given *g*_2_(*τ*) curve, the 2-pulse modulated 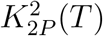 can be simulated according to Eq. 7. For the same *g*_2_(*τ*) curve, the corresponding traditional *K*^2^(*T*) without modulation can be simulated according to Eq. 3.

Particularly, when *g*_2_(*τ*) assumes the following form,

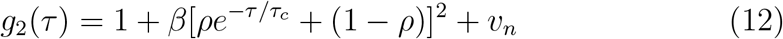

the analytical solution of the corresponding speckle contrast in the 2-pulse modulation without any approximation would be

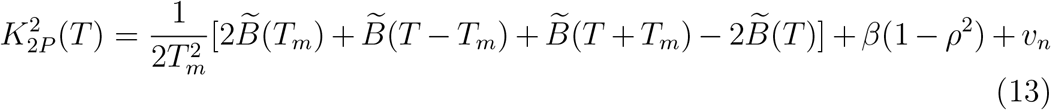

where 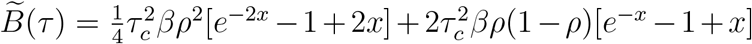 and *x* = *τ/τ*_*c*_. Similarly, plugging the *g*_2_(*τ*) model into Eq. 3, the speckle contrast without modulation would be

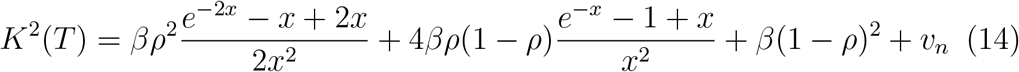

where *x* = *T/τ*_*c*_.

In simulation results presented in Fig. 2 and S5, *g*_2_(*τ*) assumes the form of Eq. 12, the parameters in which are *β* = 1, *ρ* = 1, *v*_*n*_ = 0. *T* or *τ* ranges from 10 *μ*s to 0.1 s with a resolution of 1 *μ*s. The correlation time *τ*_*c*_ is varied in simulation.

### 4.3 Instrumentation setup

A volume holographic grating (VHG) wavelength stabilized laser diode (785 nm; LD785-SEV300, Thorlabs) is used to provide the light source. An optical isolator (Electro-Optics Technology, Inc.) based on Faraday rotation effect is placed immediately after the laser output to prevent potential inadvertent back reflections from disrupting the laser source. The light passing through the isolator is then coupled into a single-mode fiber (P3-780A-FC-2, Thorlabs, Inc.) to reshape the beam profile into a circular Gaussian one. The output beam is sent into an acoustic optical modulator (AOM) (AOMO 3100-125, Gooch & Housego, Inc.) through which the power of the 1st order diffraction can be manipulated. The 0th order diffraction is filtered out by an aperture. Apart from the widefield illumination in which the output beam from AOM is deflected down and incident upon the imaging object directly, another focused illumination beam path is constructed with two convex lenses (focal length, L3: 100 mm, L4: 50 mm) whose relative distance can be adjusted to focus the beam onto the imaging spot. For detection path, the light is collected by a Nikon 24mm camera lens (AF NIKKOR 24 mm f/2.8D, Nikon, Inc.) and split into two by a 50:50 plate beamsplitter (BSW17, Thorlabs, Inc.). The transmission split is focused on a camera (acA1920-155 um, Basler, Inc.) via a Nikon 50mm lens (AF NIKKOR 50 mm f/1.8D, Nikon, Inc.). The same model of lens is used to focus the reflection split onto a fiber coupler to which a single-mode fiber (P3-780A-FC-2, Thorlabs, Inc.) is attached. The output light of the fiber is collimated via a collimator (focal length: 11 mm; F220APC-780, Thorlabs, Inc.) and focused on the photosensitive area of the APD (APD410A, Thorlabs, Inc.) by a spherical lens (focal length: 40 mm; LBF254-040-B, Thorlabs, Inc.). The intensity signal measured by APD is filtered by a low-pass filter (500 kHz; EF506, Thorlabs) and sampled by the data acquisition board at 1 MHz (USB-6363, National Instrument, Inc.).

### 4.4 LSCI experimental validation in vitro and in vivo

A single-channel microfluidics device is used to test the method *in vitro*. Its bulk is manufactured with polydimethylsiloxane (Dow Corning Sylgard 184 PDMS) in a 10 to 1 base-to-curing agent mixture by weight. Titanium dioxide (CAS 1317-80-2, Sigma, USA) is added into the mixture (1.8‰ w/w) to create optical properties mimicking the tissue^40^. The scattering solution flowing through the channel is made by diluting the Latex microsphere suspensions (5100A, 10% w/w, Thermo Fisher Scientific, USA) in a 4.8% v/v ratio with distilled water to mimic the optical properties of blood.

The mouse cranial window preparation procedures were detailed by Kazmi et al.^41^. All animal procedures are approved by the Institutional Animal Care and Use Committee (IACUC) of University of Texas at Austin.

In 2-pulse modulated multiple-exposure imaging, 15 camera exposure times were used for demonstration of characterizing *g*_2_(*τ*) at multiple time lags. *T*_*m*_ = 10 *μ*s and *T* ranges from 10 *μ*s to 5 ms. Specifically, the 15 T are 10 *μ*s, 12 *μ*s, 15 *μ*s, 20 *μ*s, 30 *μ*s, 40 *μ*s, 50 *μ*s, 75 *μ*s, 100 *μ*s, 250 *μ*s, 500 *μ*s, 750 *μ*s, 1 ms, 2.5 ms and 5 ms. The raw image size is 1000 × 750. Speckle contrast is computed spatially from raw images according to Eq. 2 with a 7 × 7 sliding window. Focused illumination is employed for both APD and camera measurements. For APD measurement, the laser power is 100 mW. In camera measurements, the laser power is attenuated by AOM to avoid pixel saturation. For *in vitro* experiements, 150 raw speckle images are collected for each camera exposure time and the raw intensity signal is recorded by APD for 10 s. The measurement is repeated 5 times for each flow rate. The flow rate increases from 0 to 100 *μ*L/min with a step size of 10 *μ*L/min in each repetition. The maximum and minimum ICT values in those five repetitions are discarded and the rest three are used for the ICT comparison between camera and APD measurements. For *in vivo* measurements, 30 raw camera images are collected for each exposure time and 2 s APD signal is recorded. The measurement is repeated 5 times at each point. Data collection is performed at 28 points in cranial windows of 4 mice (C57BL/6, Charles River Laboratories Inc.).

The *g*_2_(*τ*) curve is calculated from APD recordings in software according to 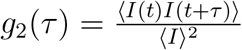 with *τ* equally spaced. The correlation time is extracted from *g*_2_(*τ*) curve by fitting to the following model

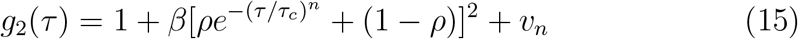

where *β* is the instrumentation factor ranging from 0 to 1, *ρ* denotes the fraction of dynamic component in the detected light ranging from 0 to 1, and *v*_*n*_ denotes the noise. *n* determines the type of *g*_1_(*τ*) model to use. *n* can be fixed to 1 or chosen from 2, 1 or 0.5 based on *R*^2^. For equally spaced *τ, g*_2_(*τ*) is concentrated in the tail when *g*_2_(*τ*) is plotted in the logarithmic *τ* scale. To counteract the skewing effects of denser *g*_1_(*τ*) sampling towards larger *τ* in the logarithmic *τ* axis, weighted fitting is deployed with 1*/τ* as the weighting function. The weighting function *w* = 1*/τ* equalizes the integral weight of data points within different *τ* ranges of the same length in the logarithmic scale, i.e. 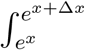 *w*(*τ*)*dτ* ∝ Δ*x* for ∀*x* ∈ ℜ. Weighted fitting by 1*/τ* improves the fitting performance in the head of *g*_2_(*τ*) curve (Supplemental Fig. S4). To match the *τ* range in 2PM-MESI, the *g*_2_(*τ*) curve is truncated in the head such that only data of *τ* ≥ 10 *μs* is used for correlation time extraction.

For both focused and widefield illumination experiments, the correlation time *τ*_*c*_ is extracted from measured 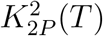 curves according to the following model

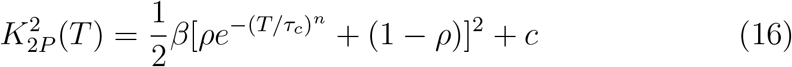

where *c* represents a constant term independent of *T*. *β, ρ* and *n* have the same meaning as those in Eq. 15. *T* is the period of the 2-pulse modulation waveform.

### 4.5 Applying 2-pulse modulation to FCS

The FCS data comes from a public FCS dataset (https://github.com/FCSlib/FCSlib/blob/master/Sample%20Data/Cy5.tif, the *g*_2_(*τ*) curve of this sample data is provided in the Figure 5.3 of its user guide). The sampling rate is 500 kHz. The *g*_2_(*τ*) curve is calculated according to 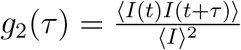 with the binned photon count number as the *I*. 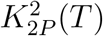 is calculated from binned photon count with the pulse width *T*_*m*_ = 2 *μ*s. Two binned photon count numbers at different time points are added together and the sum’s variance over its mean squared is calculated as 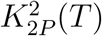 (Eq. 2). The temporal gap *T* between the two time points elongates so that 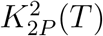 at different *T* is obtained.

## Acknowledgements

We acknowledge the support of National Institutes of Health (NIH) (Grant NS108484, EB011556) and UT Austin Portugal Program. We thank Dr. Jeanne Stachowiak, Dr. Carl C. Hayden and Dr. Feng Yuan for the help and support in the FCS project.

## Author Contributions

Q. F. and A. K. D. proposed the idea and developed the theory. Q. F. designed the experiments, did the numerical simulation, *in vitro* and *in vivo* LSCI experiments. A. T performed mouse relevant operations: surgery, handling, anesthesia. Q. F., A. T. and A. K. D. wrote the manuscript together.

## Data Availability

The sample FCS data and MATLAB processing scripts relevant to Fig. 5 can be accessed through Github (the 2PM-FCS project). URL: https://github.com/2010511951/2PM-FCS. Other experimental data and resources will be made available upon reasonable request.

## Supplementary Material

### S1. Relating K^2^(T) and g_2_(τ) in arbitrary modulation

Define the AOM modulation function as *m*(*t*), the intact speckle signal as *I*(*t*), and the modulated speckle signal as *I*_*m*_(*t*) such that

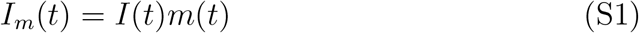

Then the intensity of pixel *i* on the camera sensor within intensity-modulated exposure time *T* would be

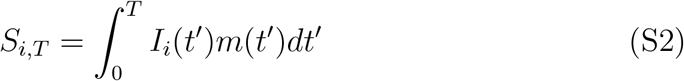

where *I*_*i*_(*t*) is the intact speckle signal of pixel *i* and *m*(*t*) is the modulation function on the illumination intensity.The second moment of modulated pixel intensity would be

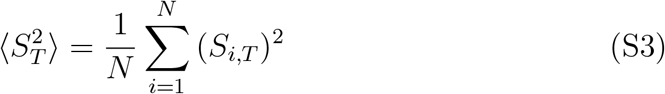

where ⟨ ⟩ denotes averaging and *N* is the number of averaged pixels. The last material needed for the derivation is the definition of intensity autocorrelation function *g*_2_(*τ*) given by Eq. S4

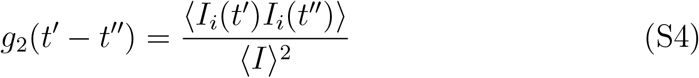

where ⟨ *I* ⟩ is the average intensity of the intact speckle signal.

Based on Eq. S1 to S4, we can derive the expression of the second moment of modulated pixel intensity with respect to the intensity modulation function *m*(*t*) and the intensity autocorrelation function *g*_2_(*τ*) of the intact signal as follows:

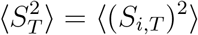

⟨ ⟩ denotes averaging over independent instances

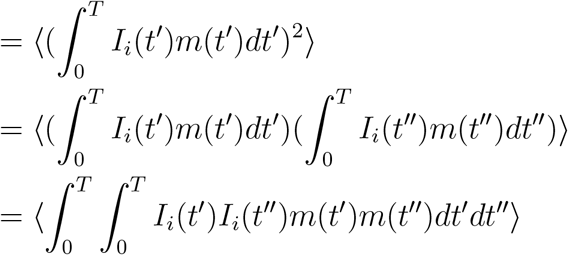

*m*(*t*) is independent of *i*

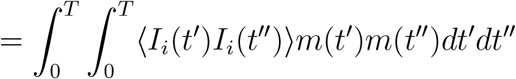

Using Eq. S4

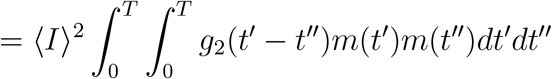

Symmetry of *t*′ and *t*′′; *g*_2_(*τ*) is even

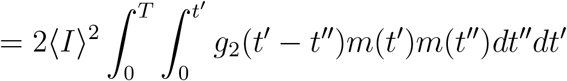

Let *t*′ − *t*′′ = *τ, then t*′′ = *t*′ − *τ, dt*′′ = −*dτ*

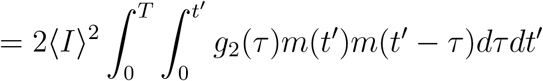

Change the order of integral

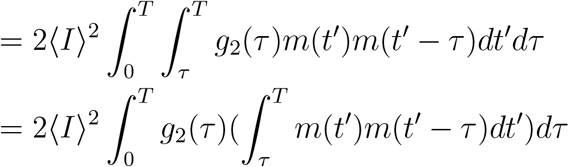

Let *t* = *t*′ − *τ*, then *t*′ = *t* + *τ, dt*′ = *dt*

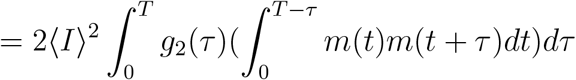

Define

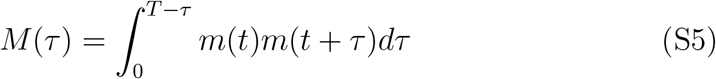

then

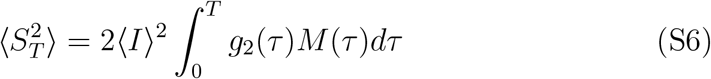

Since

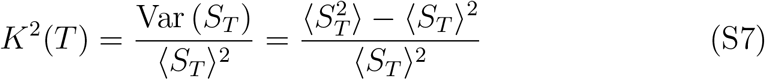

where ⟨ *S*_*T*_ ⟩ is the mean pixel intensity of modulated speckle signal within exposure time *T* and ⟨ *S*_*T*_⟩ = *T* ⟨ *I*_*m*_⟩ where ⟨ *I*_*m*_⟩ is the mean intensity of the modulated speckle signal, we arrive at the expression of speckle contrast of the within-exposure modulated speckle signal (Eq. S8).

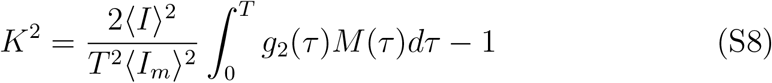

Notice that when the modulation function *m*(*t*) is a constant 1, we have *M* = *T* − *τ* and Eq. S8 reduces to the expression of speckle contrast that is commonly seen (Eq. 3). In other words, the classic expression of speckle contrast we use is a particular case of Eq. S8 when the illumination intensity is held constant. Finally, we would like to introduce one important observation about *M* (*τ*) (Lemma S1.1).

#### Lemma S1.1

(Integral property of *M* (*τ*))

*If the average intensity of the intact speckle signal I*(*t*) *remains steady over time, i*.*e*., 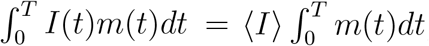, *then the integral of M* (*τ*) *satisfies* 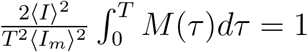.

*Proof*. Because *I*_*m*_(*t*) = *I*(*t*)*m*(*t*), we have

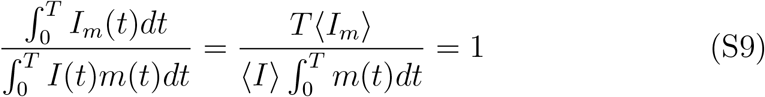

Hence,

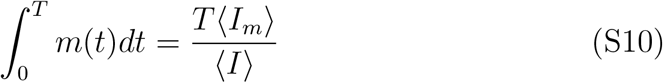

Therefore,

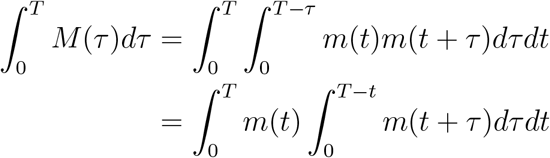

Let *t* = *t*′ = *t + τ*, then *dt*′ *= dτ*

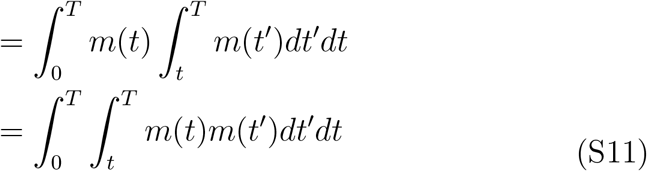

Symmetry of *m*(*t*) and m(*t*′)

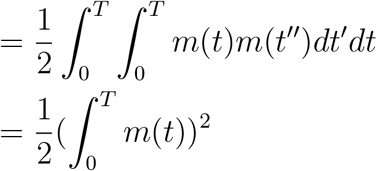

Plug in Eq. S10

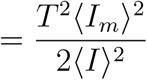

Namely, 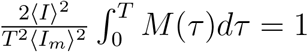. The proof is over. □

### S2. 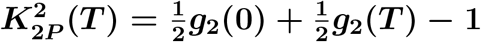 if m(t) = δ(0) + δ(T)

*Proof*. Denote *I*(*t*) as *I* and *I*(*t*+*τ*) as *I*_*τ*_, then according to 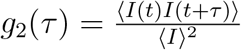
we have

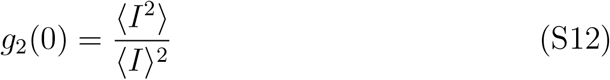

and

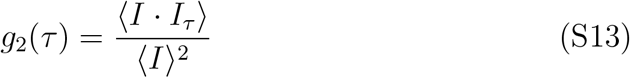

Since Var (*I*) = ⟨ *I*^2^⟩ −⟨ *I*⟩ ^2^ and Cov (*I, I*_*τ*_) = ⟨ *I I*_*τ*_⟩ *−* ⟨ *I*⟩^2^ where Var (*X*) and Cov (*X, Y*) denote the variance of *X*, and the covariance between *X* and *Y*, we have

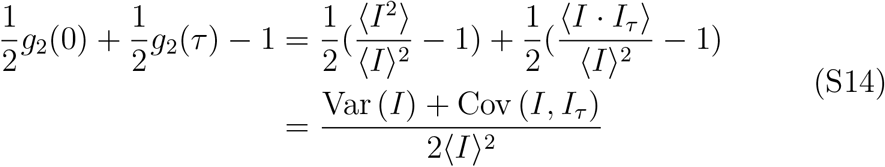

If *m*(*t*) = *δ*(0) + *δ*(*τ*), the pixel intensity *S* would be *S* = *I* + *I*_*τ*_ and 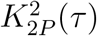 would be

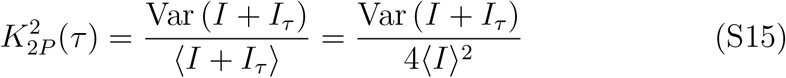

Therefore, to prove that 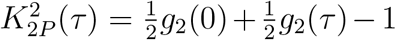, based on Eq. S14 and

S15, one only needs to prove that Var (*I* + *I*_*τ*_) = 2 Var (*I*) + 2 Cov (*I, I*_*τ*_), which is true since Var (*I* + *I*_*τ*_) = Var (*I*) + Var (*I*_*τ*_) + 2 Cov (*I, I*_*τ*_) and Var (*I*) = Var (*I*_*τ*_). The proof is over. □

### S3. The Impact of Non-zero Residual Illumination

We can model the non-zero residual illumination in 2-pulse modulation as

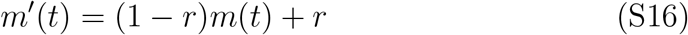

where *r* is the relative amplitude of residual illumination during the off state and ranges from 0 to 1. *m*(*t*) here is the ideal 2-pulse modulation with zero residual illumination, and ranges between 0 and 1. Then the modulation autocorrelation function would be

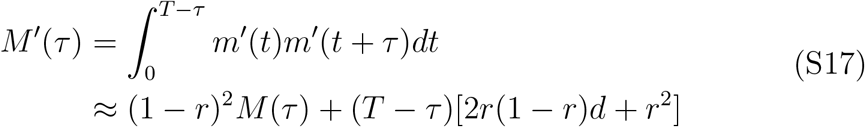

where 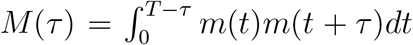 and *d* is the duty cycle of *m*(*t*) or the pseudo duty cycle of *m*^′^(*t*). Fig. S1a shows an example of how *M* (*τ*) would be skewed in presence of a non-zero residual illumination (r=0.1). The square of speckle contrast corresponding to *m*^′^(*t*) would then become

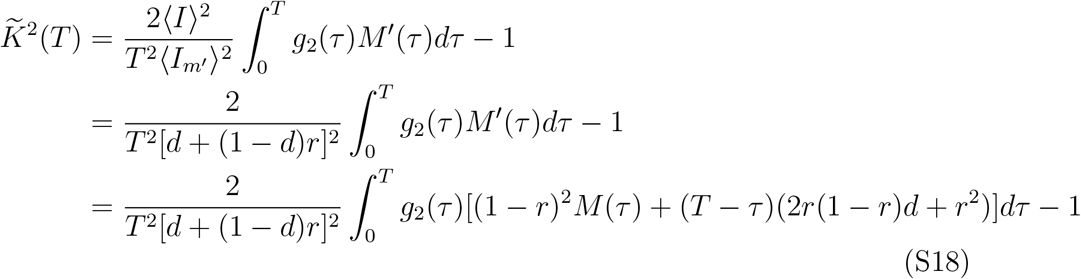

Simplify Eq. S18, we get

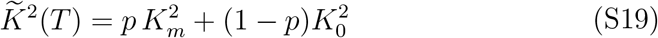

where 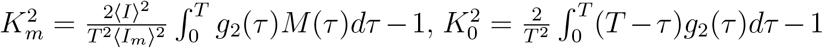, and 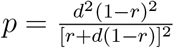. Therefore, the square of speckle contrast, *K*^2^ in presence of a non-zero residual illumination in 2-pulse modulation would be the weighted sum of that of an ideal 2-pulse modulation plus that of no modulation on intensity. *p* indicates the proportion of the contribution by the ideal 2-pulse modulation. It is noticed that when *r* increases, *p* drops and that when *d* increases, *p* rises. Fig. S1b shows an example of how an AOM with limited OD when gating the light would affect the tail of 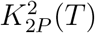 curves when *T* is large.

### S4. The impact of pulse duration on the accuracy of measuring absolute and relative values of *g*_*2*_(*τ*)

In this section, we would like to answer the question of how to choose the pulse duration when doing 2-pulse modulated multiple exposure imaging. We demonstrated the validity of a 10 *μ*s pulse duration in extracting correlation times as short as 30 *μ*s (Fig. 3f). But it does not have to be always the case. The pulse duration can be longer when measuring *g*_2_(*τ*) of slowly varying signals. We examined the optimal pulse duration selection through numerical simulation. For a given pulse duration *T*_*m*_, we evaluated the discrepancy between *g*_2_(*τ*) and its estimation by 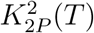 at various correlation times (Fig. S5). For a given pulse duration, the maximum percent discrepancy between 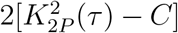and the absolute value of *g*_2_(*τ*) decreases as *τ*_*c*_ increases (Fig. S5a). When *τ*_*c*_ becomes larger than 10 times *T*_*m*_, the percentage discrepancy drops below 0.2%. In other words, to recover the absolute value of *g*_2_(*τ*) of the signal of interest within a maximum of 0.2% discrepancy threshold, the pulse duration *T*_*m*_ should be made shorter than 10% of the correlation time *τ*_*c*_ of the signal. On the other hand, if the correlation time is the only interest about *g*_2_(*τ*), i.e., the relative value of *g*_2_(*τ*) or 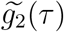 is of interest, then the pulse duration can be longer than 10% of *τ*_*c*_ (Fig. S5b). But considering that 2-pulse modulated multiple exposure imaging can only capture *g*_2_(*τ*)’s shape in the range of *τ* ≥ *T*_*m*_, it is recommended that *T*_*m*_ not be longer than *τ*_*c*_ to ensure sufficient sampling of the exponential-decay phase of *g*_2_(*τ*).

**Figure S1:**
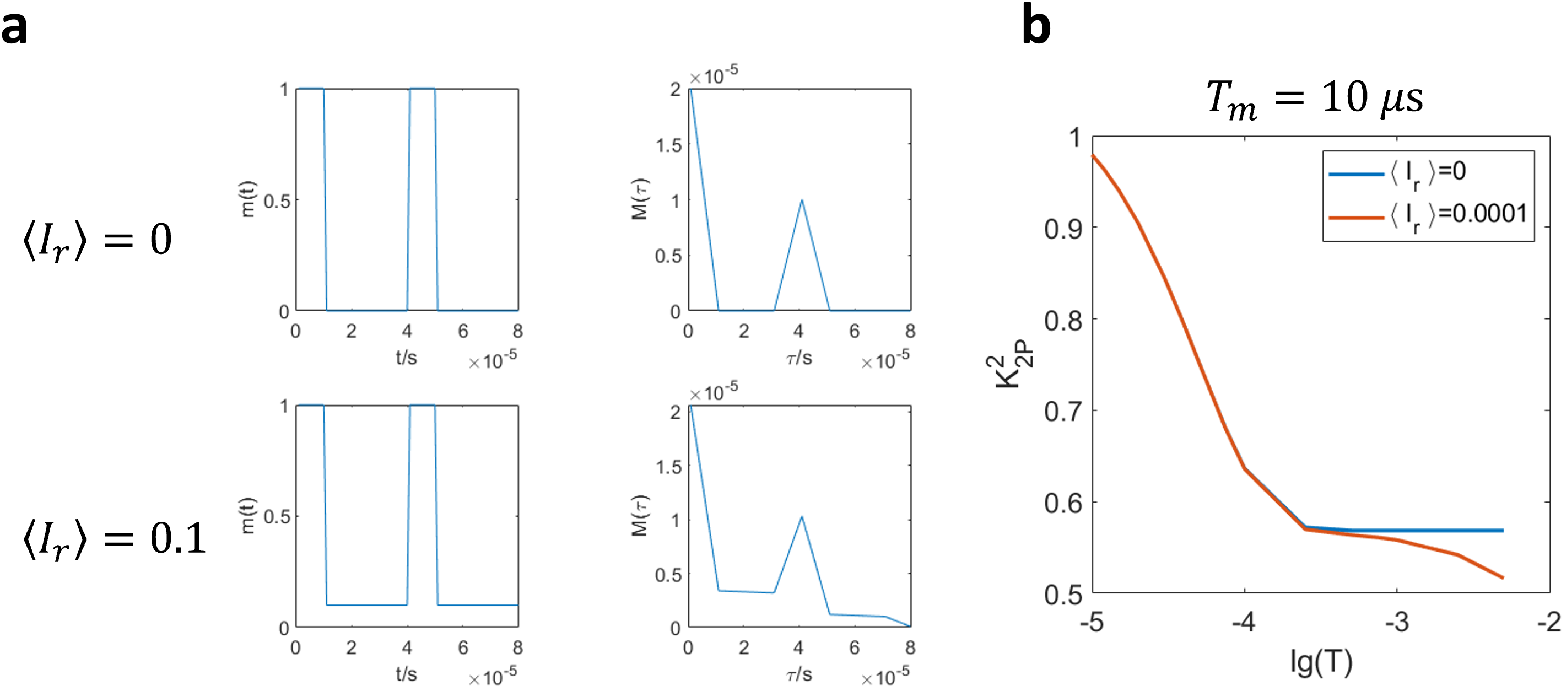
The impact of non-zero residual illumination between two illumination pulses 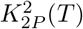. **a** How the modulation autocorrelation function *M* (*τ*) would be skewed by a non-zero residual illumination (r=0.1). **b** The comparison of 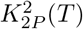 curves with and without residual illumination. An AOM with an OD of 4 when gating the light is simulated for the former case.

**Figure S2:**
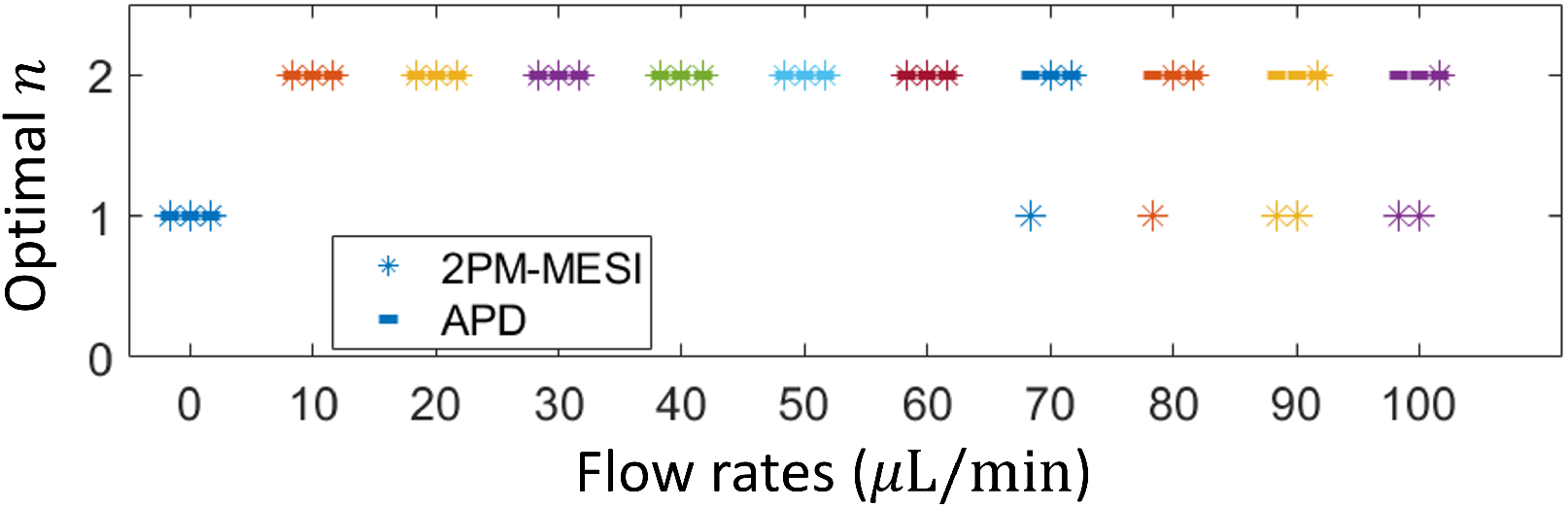
The optimal *n* given by the fitting algorithm in various flow rates. Dashed line: APD. Asteroids: 2PM-MESI. For each flow rate, the experiment is repeated for five times. Three of the five repeats are shown here and grouped together by the same color in the plot. Different colors represent different flow rates. When the flow rate is zero, the optimal *n* is 1, which is true for both APD and 2PM-MESI fitting results. When the flow rate is not zero, the optimal *n* is 2 according to APD fitting results. 2PM-MESI identifies the same optimal *n* for small flow rates (≤ 60 *μ*L/min). But for higher flow rates, instability in estimating the optimal *n* is observed, which could be due to the downticking tail of the 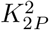 curve induced by the non-zero residual illumination between illumination pulses.

**Figure S3:**
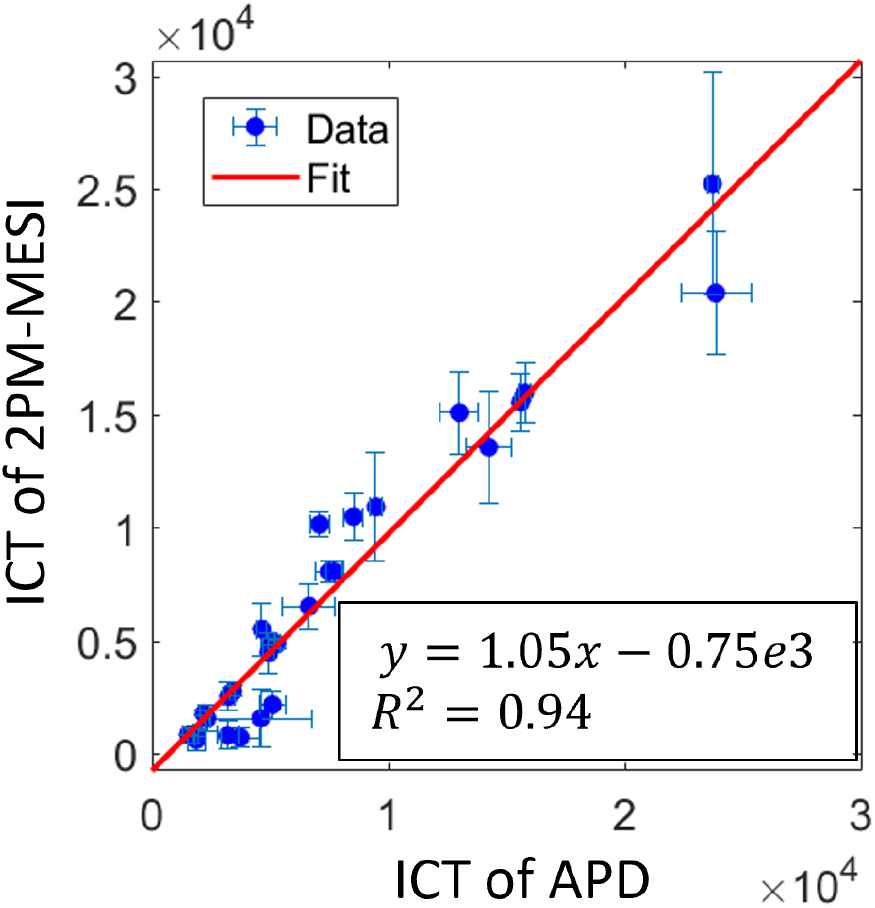
Comparison of ICT values extracted from *g*_2_(*τ*) and 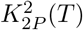 with unfixed *n*. 28 points from 4 mice.

**Figure S4:**
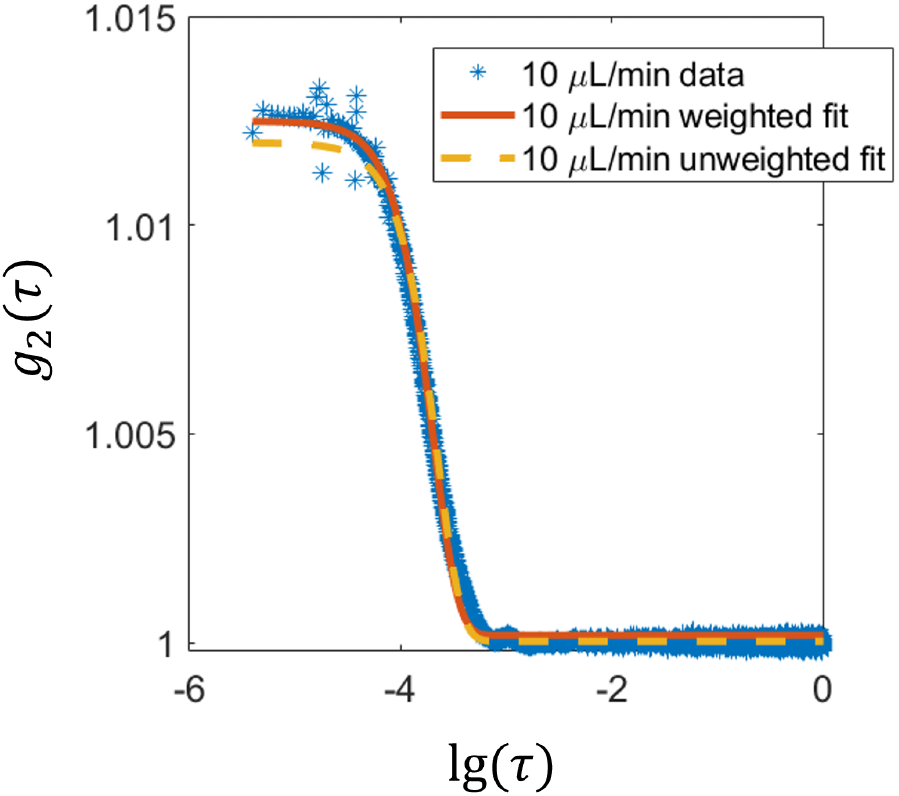
Comparison of the performance of weighted fitting vs. unweighted fitting. The weighted fitting by 1*/τ* improves the fitting performance in the head of *g*_2_(*τ*) curve compared with unweighted fitting.

**Figure S5:**
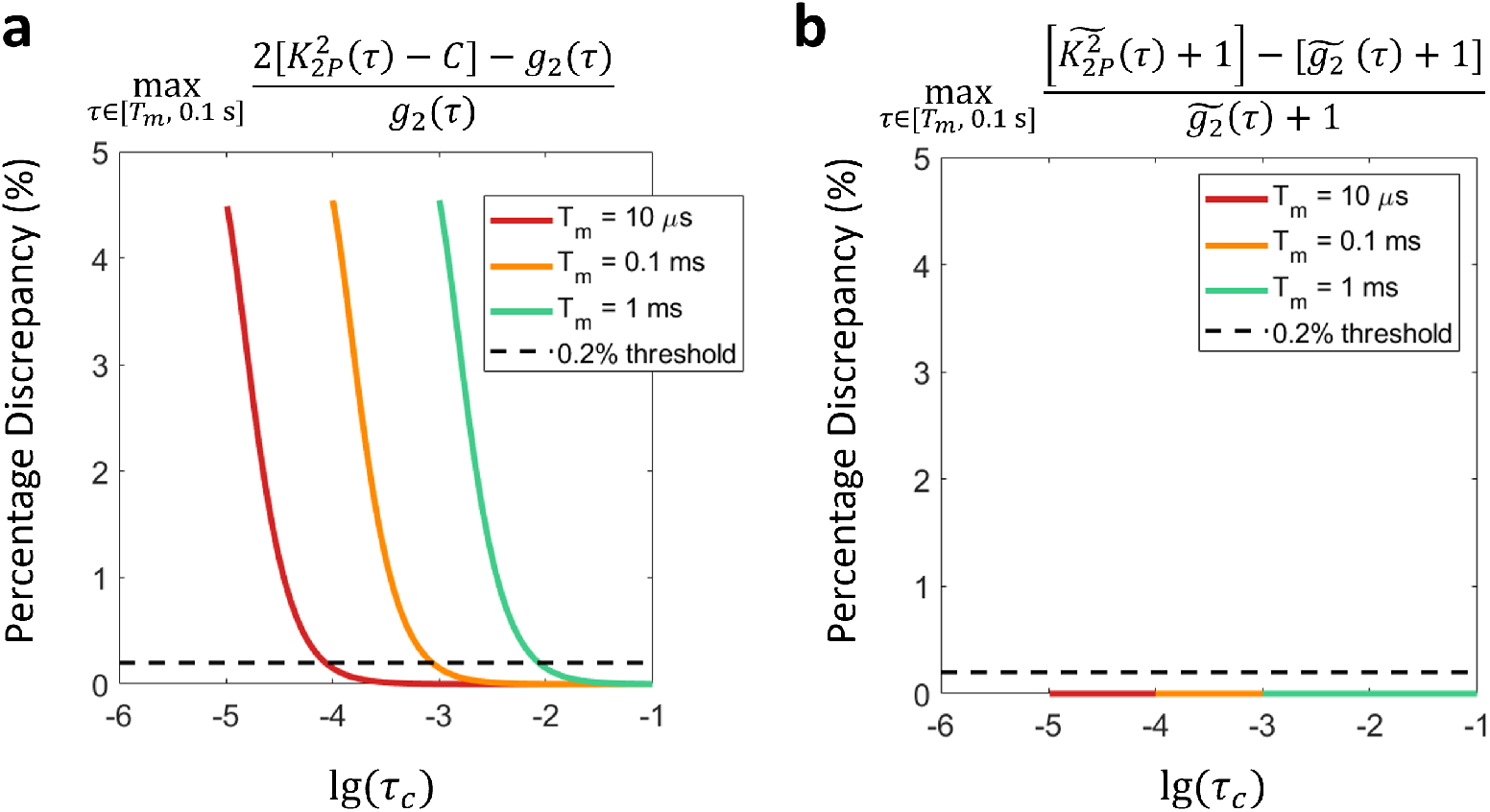
The accuracy of estimating *g*_2_(*τ*) and 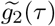 based on 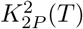 for signals of different correlation times. **a** The maximum percentage discrepancy between absolute *g*_*2*_ (*τ*) and that estimated by 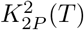. The *y*-axis is max 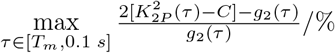. **b** The maximum percentage discrepancy between normalized *g*_2_(*τ*) and 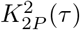. The *y*-axis is max 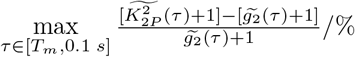.

